## Supplementary Figures for "Disentangling the mutational effects on protein stability and interaction of human MLH1"

### *Supplementary Information*


**Fig. S1**, *AlphaFold2-Multimer prediction of the C-terminal MutLα complex.* *p.2*

**Fig. S2**, *Vector map of pDEST-DHFR-PCA.* *p.3*

**Fig. S3**, *Vector maps of pDEST22 and pDEST32.* *p.4*

**Fig. S4**, *Distributions of abundance and interaction scores per tile.* *p.5*

**Fig. S5**, *Score correlation between replicates.* *p.6*

**Fig. S6**, *Heatmaps of the standard error (SE) of abundance and interaction scores.* *p.7*

**Fig. S7**, *Median score correlation with relative solvent accessible surface area (rSASA).* *p.8*

**Fig. S8**, *Validation of loss- and gain-of-interaction variants.* *p.9*

**Fig. S9**, *Interfaces of the MutLα complex.* *p.10*

**Fig. S10**, *Prediction of a novel interaction site with PMS2.* *p.11*

**Fig. S11**, *Correlation of Rosetta and GEMME predictions with experimental scores.* *p.13*

**Fig. S12**, *Predicting abundance and interaction scores using Rosetta and GEMME scores.* *p.14*

**Fig. S13**, *Separation of interaction scores using GEMME predictions and abundance scores.* *p.15*

**Fig. S14**, *Classification of MLH1 pathogenicity using experimental and computational scores.* *p.16*

**Fig. S15**, *Hydrophobicity of the canonical PMS2 interface.* *p.17*

**Fig. S16**, *Dynamic range and sensitivity of the DHFR-PCA and Y2H assay.* *p.18*

**Fig. S17**, *Benchmarking abundance and interaction scores against existing literature.* *p.19*

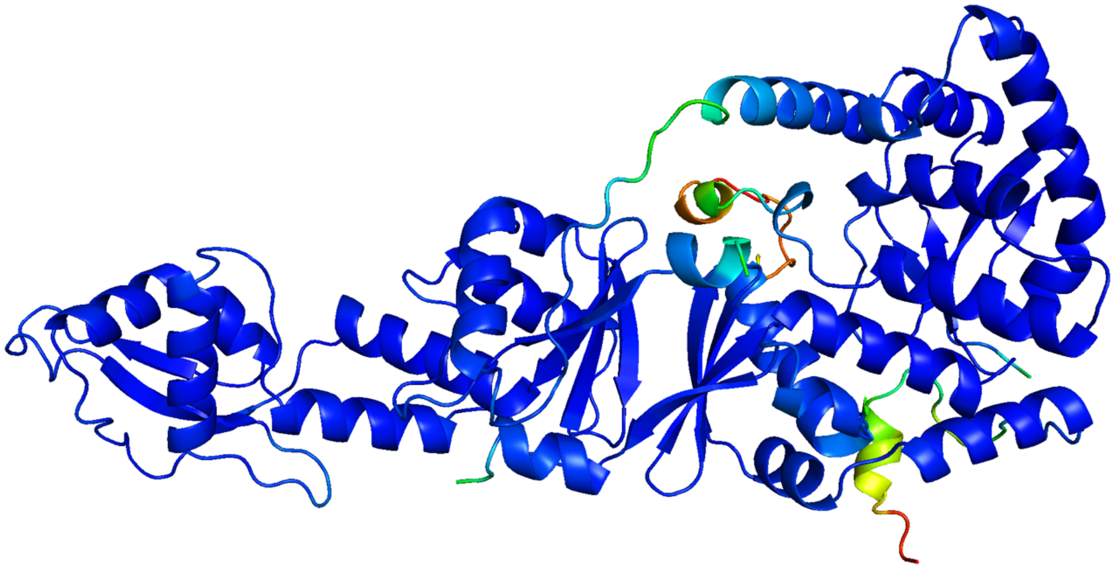

**Fig. S1** – *AlphaFold2-Multimer prediction of the C-terminal MutLα complex.* High confidence model of the human C-terminal MutLα complex predicted by AlphaFold2-Multimer. The structure is colored by the pLDDT score using the reversed rainbow color palette in PyMOL.

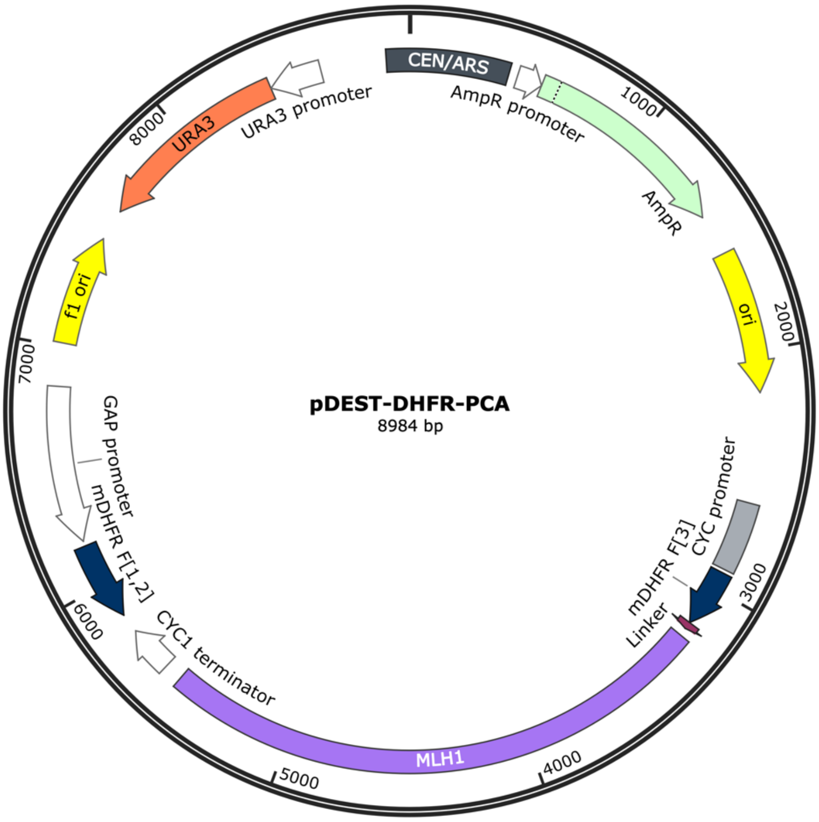


**Fig. S2** – *Vector map of pDEST-DHFR-PCA.* Vector map of the CEN-based expression vector pDEST-DHFR-PCA with a *URA3* marker. The mDHFR[F3]-MLH1 fusion protein and mDHFR[F1,2] were expressed from the CYC1 and GAP promoter, respectively. The vector map was generated using SnapGene®.

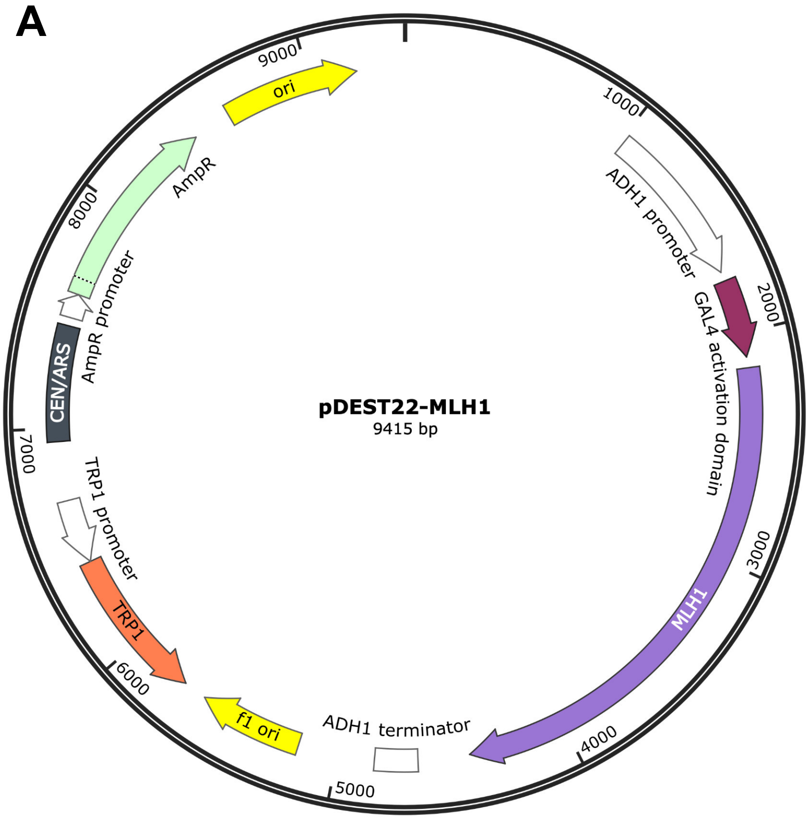


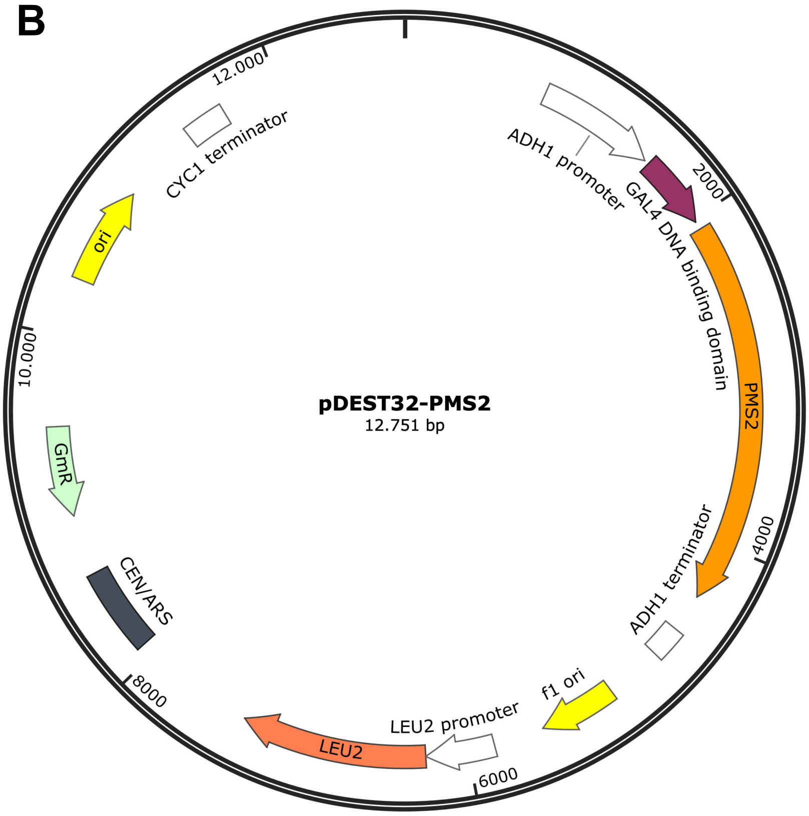


**Fig. S3** – *Vector maps of pDEST22 and pDEST32.* Vector maps of the CEN-based expression vectors pDEST22 and pDEST32 with a *TRP1* and *LEU2* marker, respectively. MLH1 was fused to the activation domain and PMS2 was fused to the DNA binding domain. The vector maps were generated using SnapGene®.

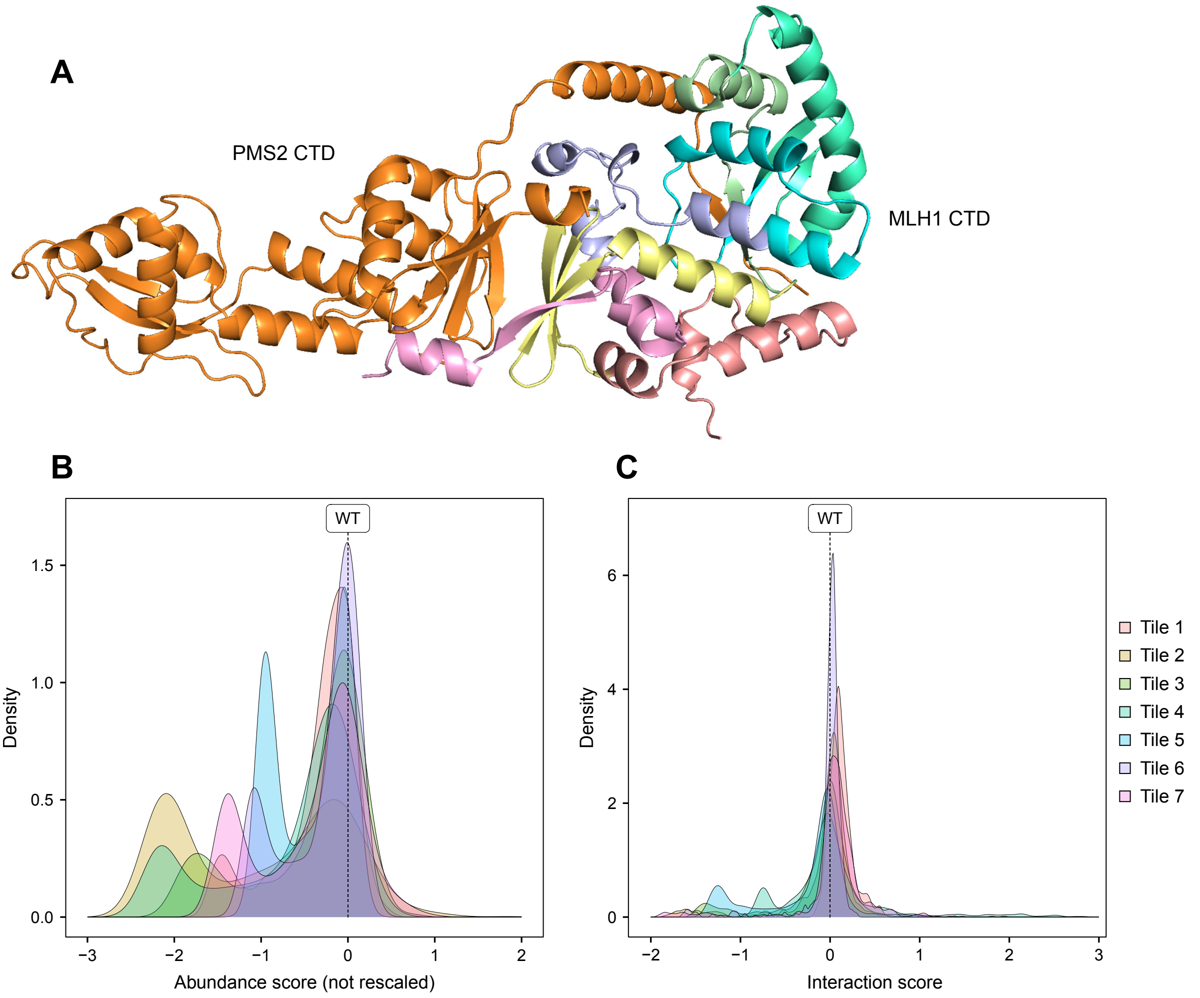


**Fig. S4** – *Distributions of abundance and interaction scores per tile.* (A) The predicted structure of the C-terminal MutLα complex with the C-terminal domain (CTD) of MLH1 colored by the seven tiles. The tiles are colored according to the legend in (C). The C-terminal domain of PMS2 is shown in orange. (B) Density plot showing the distribution of abundance scores for each tile in the C-terminal domain of MLH1 prior to rescaling. The tiles are colored according to the legend in (C). (C) Same as in (B), but for interaction scores.

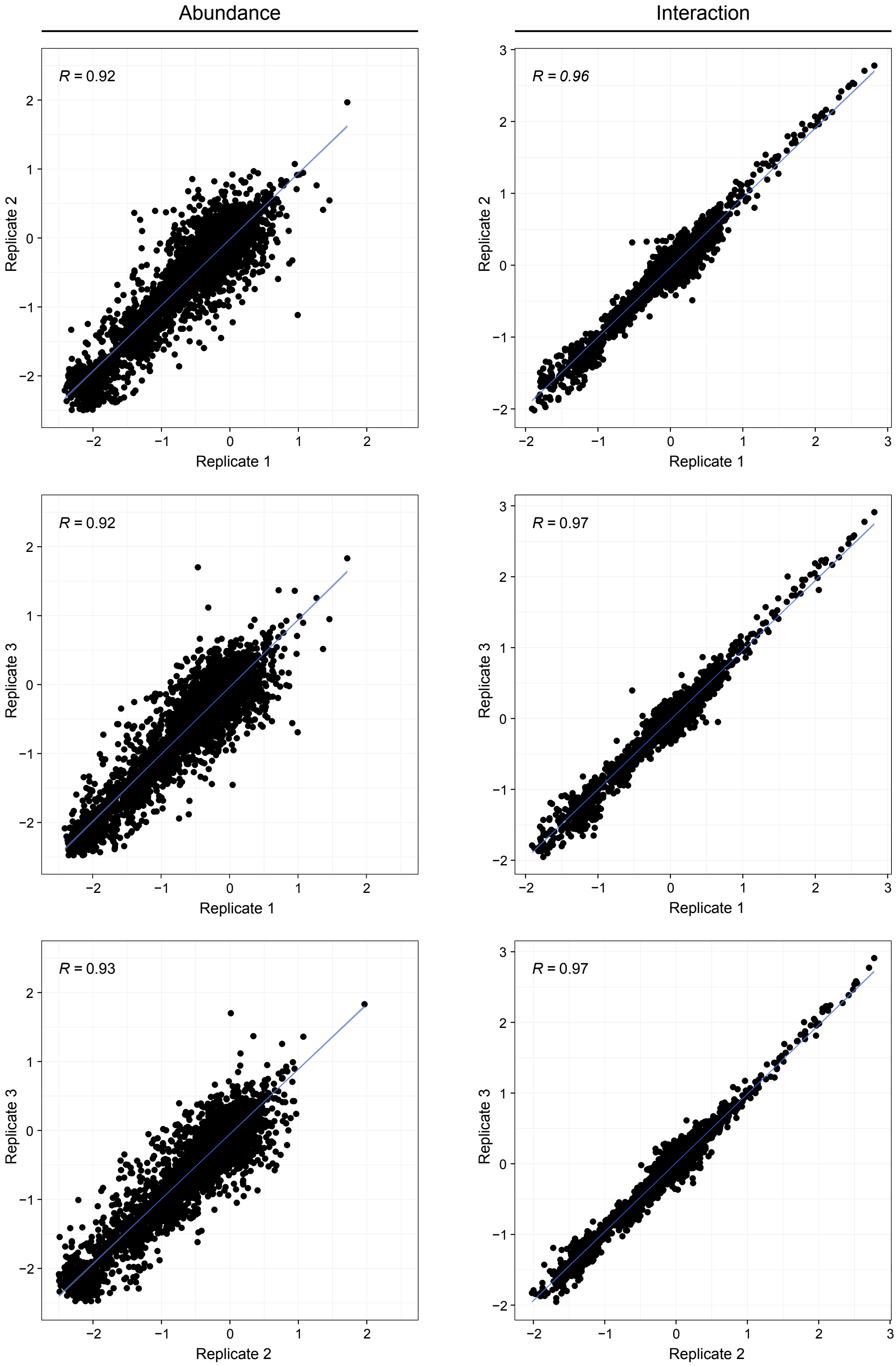


**Fig. S5** – *Score correlation between replicates*. Scatter plots showing the score correlation between the three replicates for DHFR-PCA (abundance) and Y2H (interaction). The Pearson correlation coefficient (R) is shown for each plot.

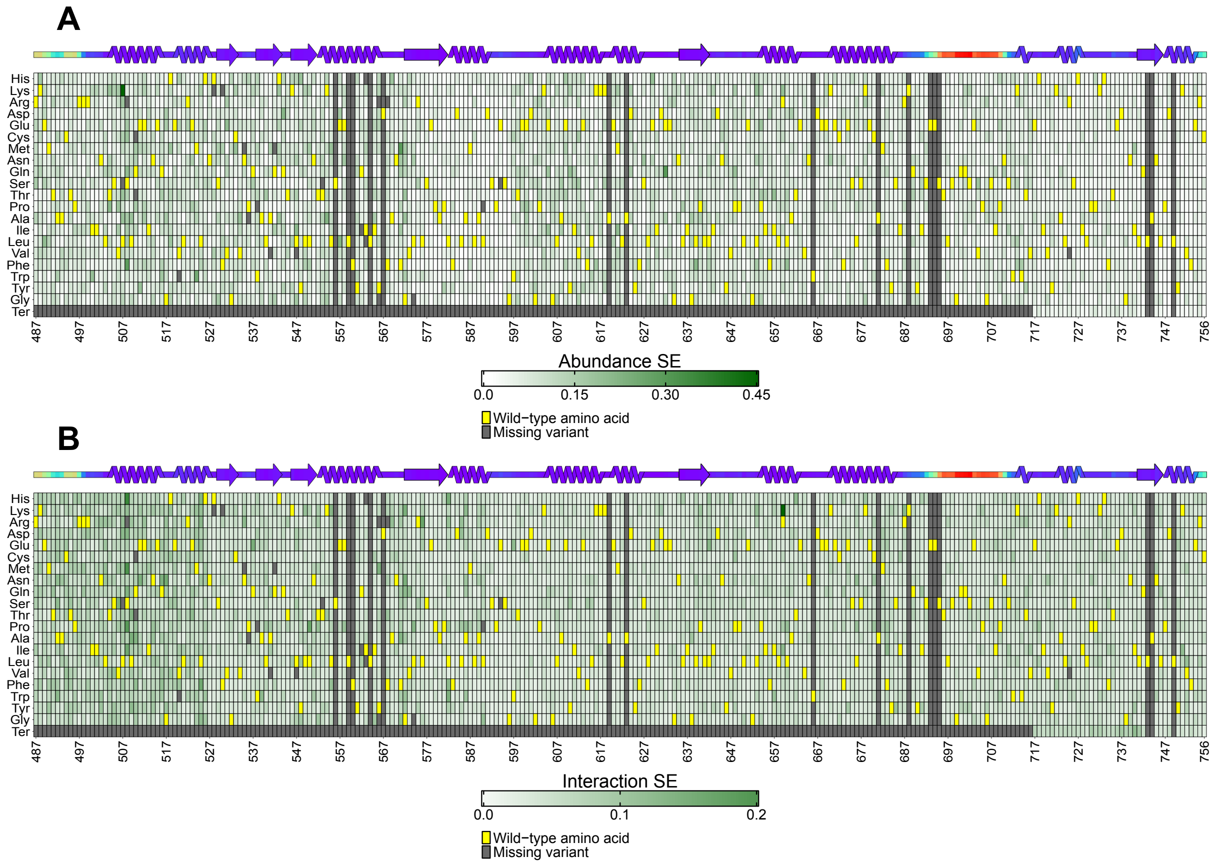


**Fig. S6** – *Heatmaps of the standard error (SE) of abundance and interaction scores.* (A) Heatmaps of the SE for the abundance scores, calculated using Enrich2. The SE has been rescaled in accordance with the rescaling of the abundance scores. The SE range from 0 (white) to 0.45 (dark green). The wild-type residue at each position is marked in yellow, while missing variants are marked in gray. The position in the full-length MLH1 protein is shown below the heatmap. A linear representation of the secondary structure of MLH1 is displayed above the heatmap, colored by AlphaFold2 pLDDT as a measure of disorder. (B) Same as in (A), but for interaction scores. Note, the color scale is different from in (A).

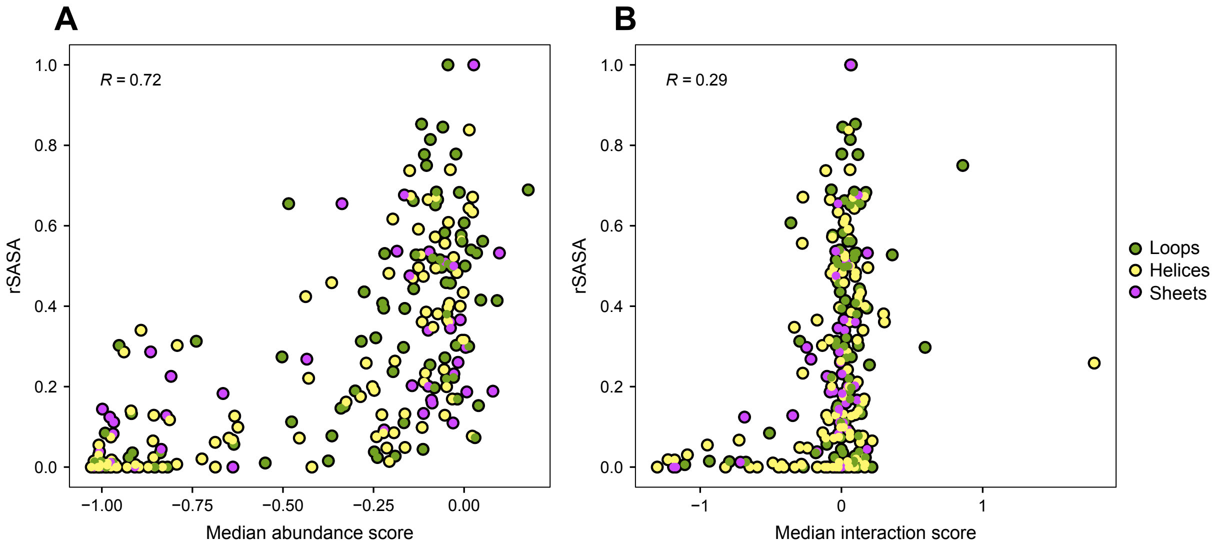


**Fig. S7** – *Median score correlation with relative solvent accessible surface area (rSASA).* (A) Scatter plot showing the rSASA plotted against the median abundance score per position. Residues are colored by the secondary structural element according to the legend in (B). A rSASA value of 0 corresponds to a completely buried residue, while 1 corresponds to a completely exposed residue. The Spearman's R correlation coefficient is shown. (B) Same as in (A), but for median interaction scores.

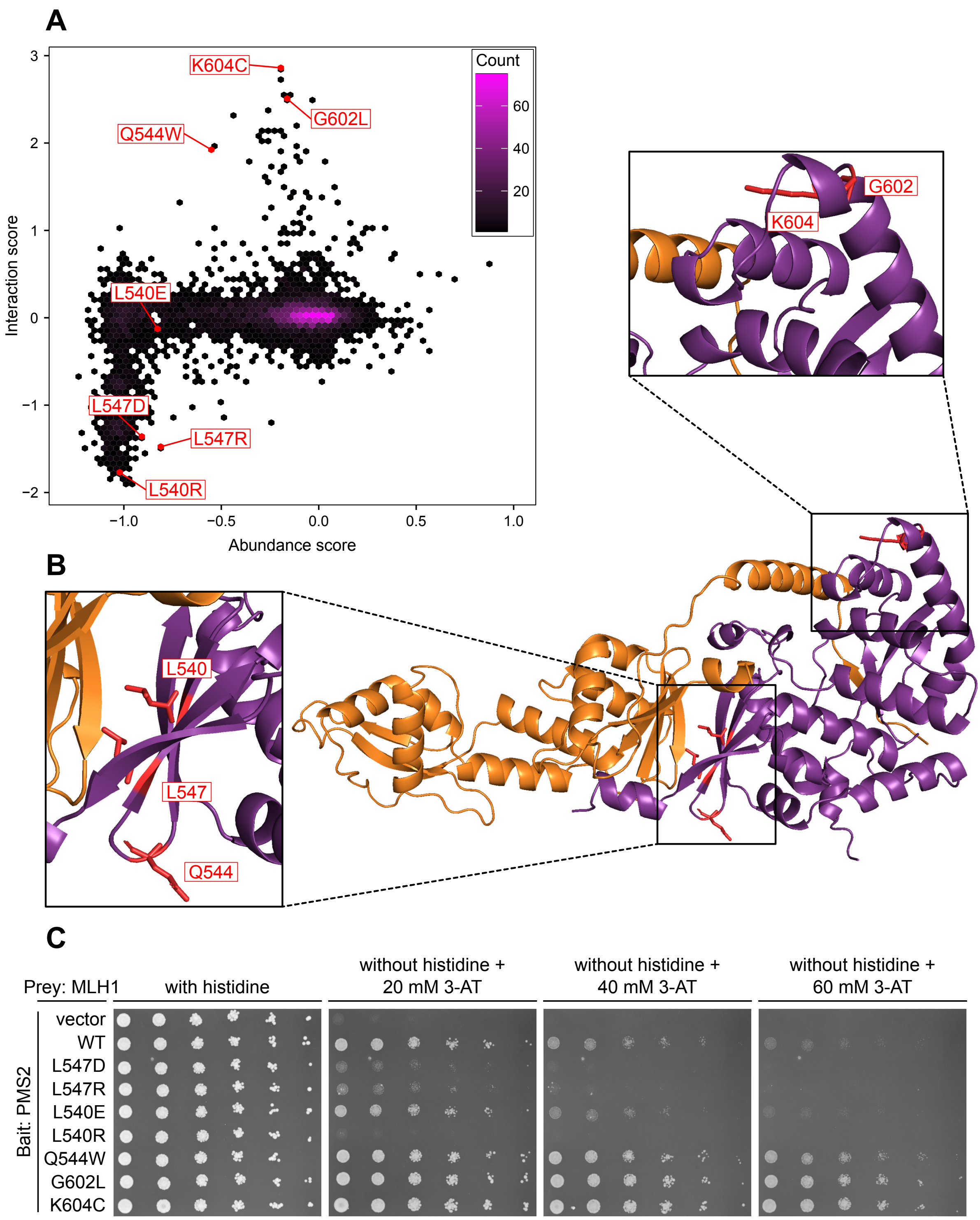


**Fig. S8** – *Validation of loss- and gain-of-interaction variants.* (A) Scatter density plot showing the interaction score plotted against the abundance score. The color scale is indicated in the legend. Selected loss-of-interaction and gain-of-interaction variants are highlighted in red. (B) The structure of the predicted C-terminal MutLα complex with the selected MLH1 variants highlighted. (C) Y2H growth assays comparing the growth of the vector, wild-type MLH1 and the indicated MLH1 variant. Cells were grown on medium with histidine and without histidine with 20 mM (the concentration used in the high-throughput screen), 40 mM or 60 mM 3-AT.

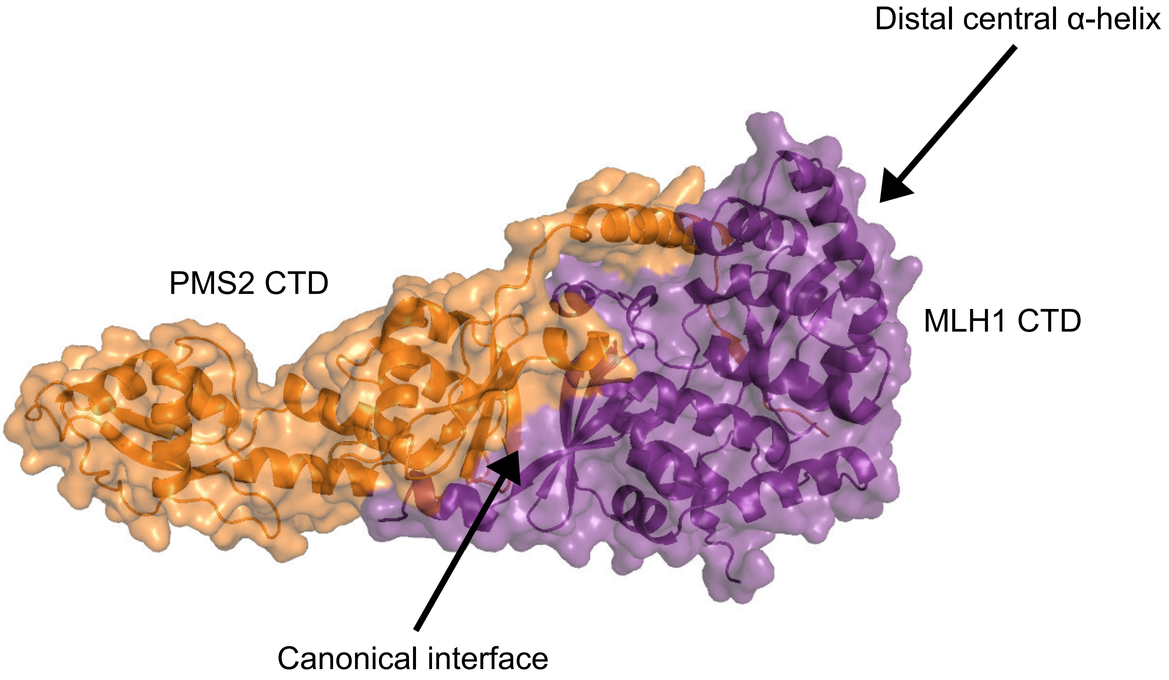


**Fig. S9** – *Interfaces of the MutLα complex.* The AlphaFold2-Multimer predicted structure of the human C-terminal MutLα complex shows the interaction between MLH1 (purple) and PMS2 (orange). The canonical interface known from the yeast complex and the novel interaction site mediated via the distal central α-helix are indicated by black arrows.

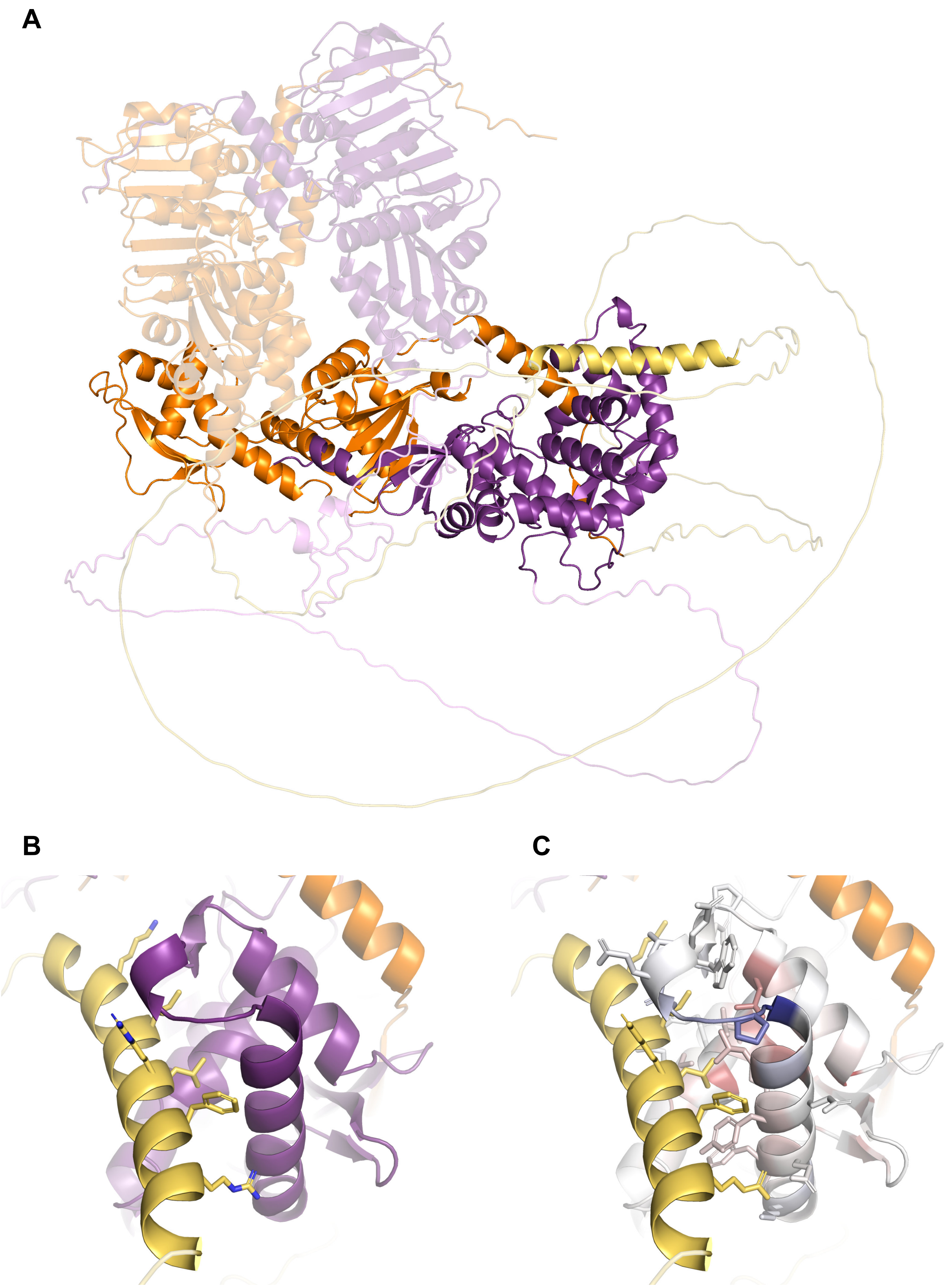


**Fig. S10 –** *Prediction of a novel interaction site with PMS2.* (A) The full structure of the human MutLα complex as predicted by AlphaFold 3. As in other figures, MLH1 is colored purple, and PMS2 is colored orange. The intrinsically disordered linker regions between the N- and C-terminal domains are colored magenta and yellow, respectively. The N-terminal domains and linker regions are displayed as transparent. AlphaFold3 predicts an α-helix within the disordered linker of PMS2, spanning residues 409-431 (yellow), which binds to the distal central α-helix in MLH1. (B) A zoomed-in view of the interaction between the α-helix in PMS2 (yellow) and the central α-helix in MLH1 (purple). Residues facing MLH1 in the PMS2 α-helix are displayed as sticks, with nitrogen atoms in sidechains highlighted in blue. (C) The same view as in (B), with MLH1 colored by the per-position median interaction score. The color scale corresponds to that shown in Fig. 3C. Contact residues in MLH1 within 6 Å of the PMS2 α-helix are shown as sticks. Notably, many of these residues locate to positions where our experiments showed both loss- and gain-of-interaction variants. For visual clarity, the coloring of nitrogen and oxygen atoms in the sidechains has been omitted.

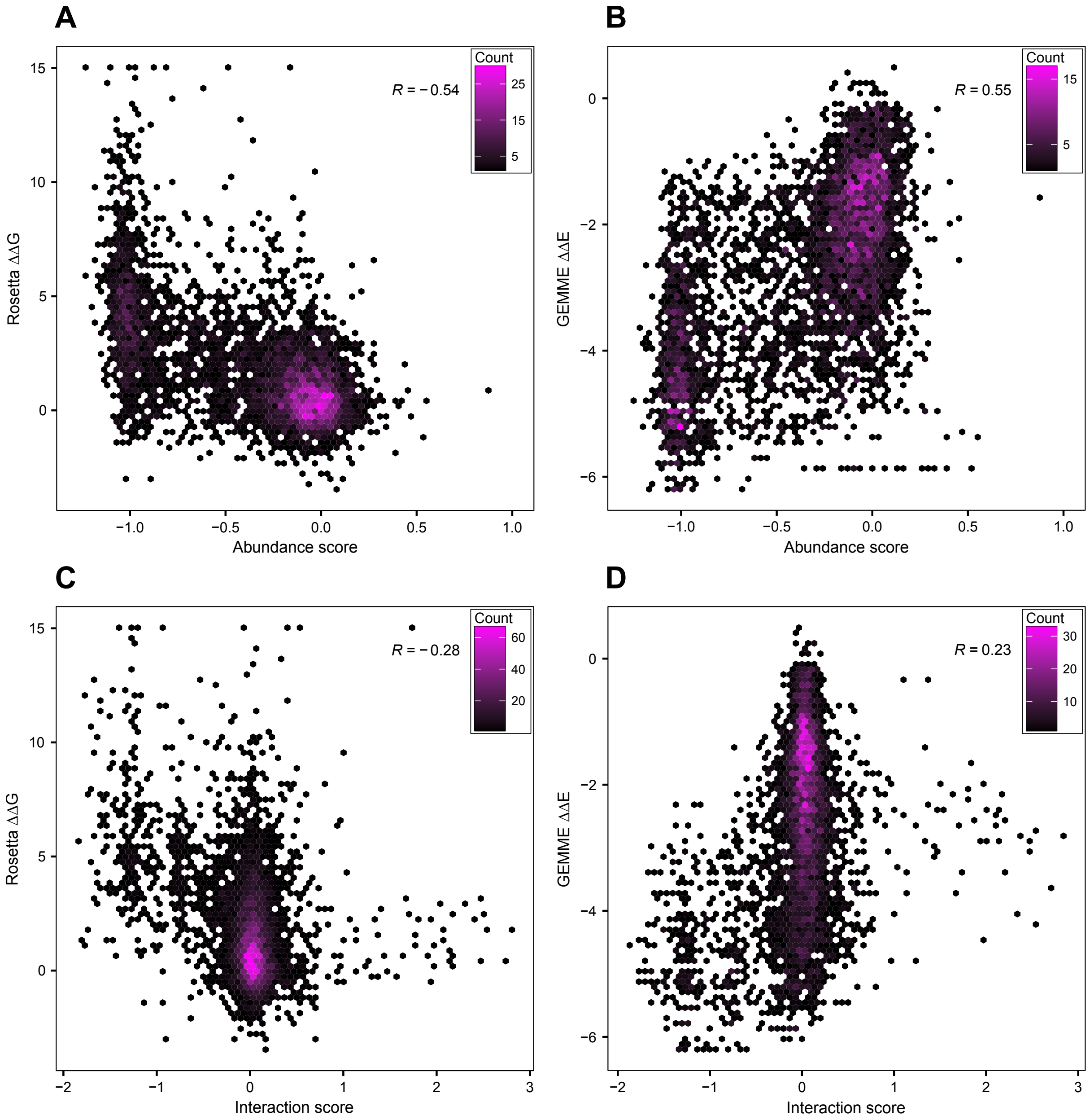


**Fig. S11** – *Correlation of Rosetta and GEMME predictions with experimental scores.* (A) Scatter density plot comparing the predicted Rosetta ΔΔG scores and the abundance scores. The color scale is indicated in the legend. The Spearman's R correlation coefficient is shown. (B) Same as in (A), but for predicted GEMME ΔΔE scores. (C) Scatter density plot comparing the predicted Rosetta ΔΔG scores and the interaction scores. The color scale is indicated in the legend. The Spearman's R correlation coefficient is shown. (D) Same as in (C), but for predicted GEMME ΔΔE scores.

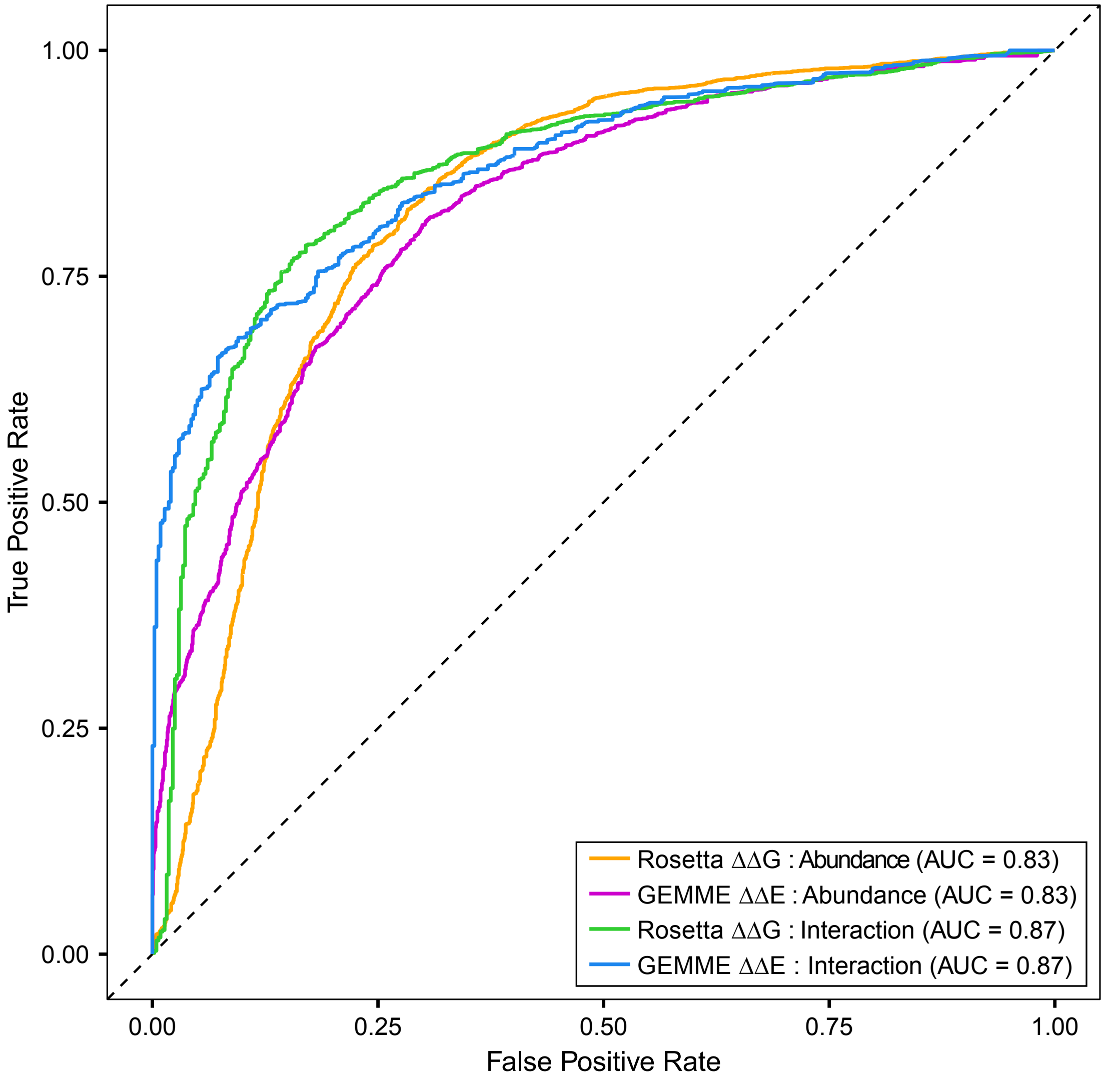


**Fig. S12** – *Predicting abundance and interaction scores using Rosetta and GEMME scores.* ROC curves to assess how well Rosetta and GEMME perform in predicting the abundance and interaction scores. A cutoff of -0.5 was applied to the abundance and interaction scores. Variants with a score below -0.5 were categorized as detrimental, while those with a score above -0.5 were categorized as wild-type-like. The AUC for each predictor is reported in the legend. The dashed diagonal line denotes performance of a random classifier. Note that variants with scores above -0.5 include both wild-type-like variants and variants with increased abundance or PMS2 interaction.

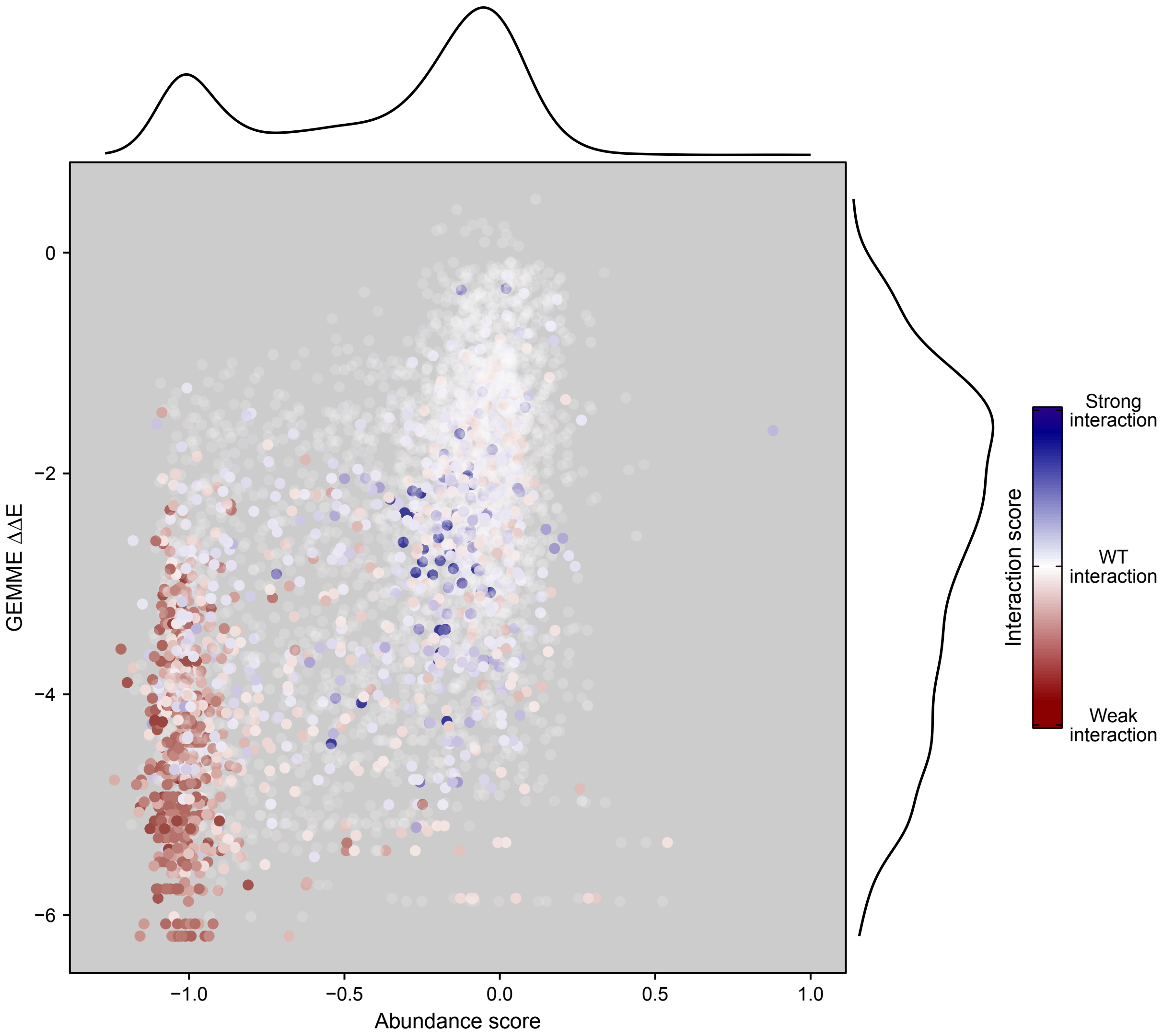


**Fig. S13** – *Separation of interaction scores using GEMME predictions and abundance scores.* Scatter plots showing the correlation between the predicted GEMME ΔΔE score and abundance scores. The MLH1 variants are colored by the interaction score and the color scale corresponds to that shown in Fig. 3A.

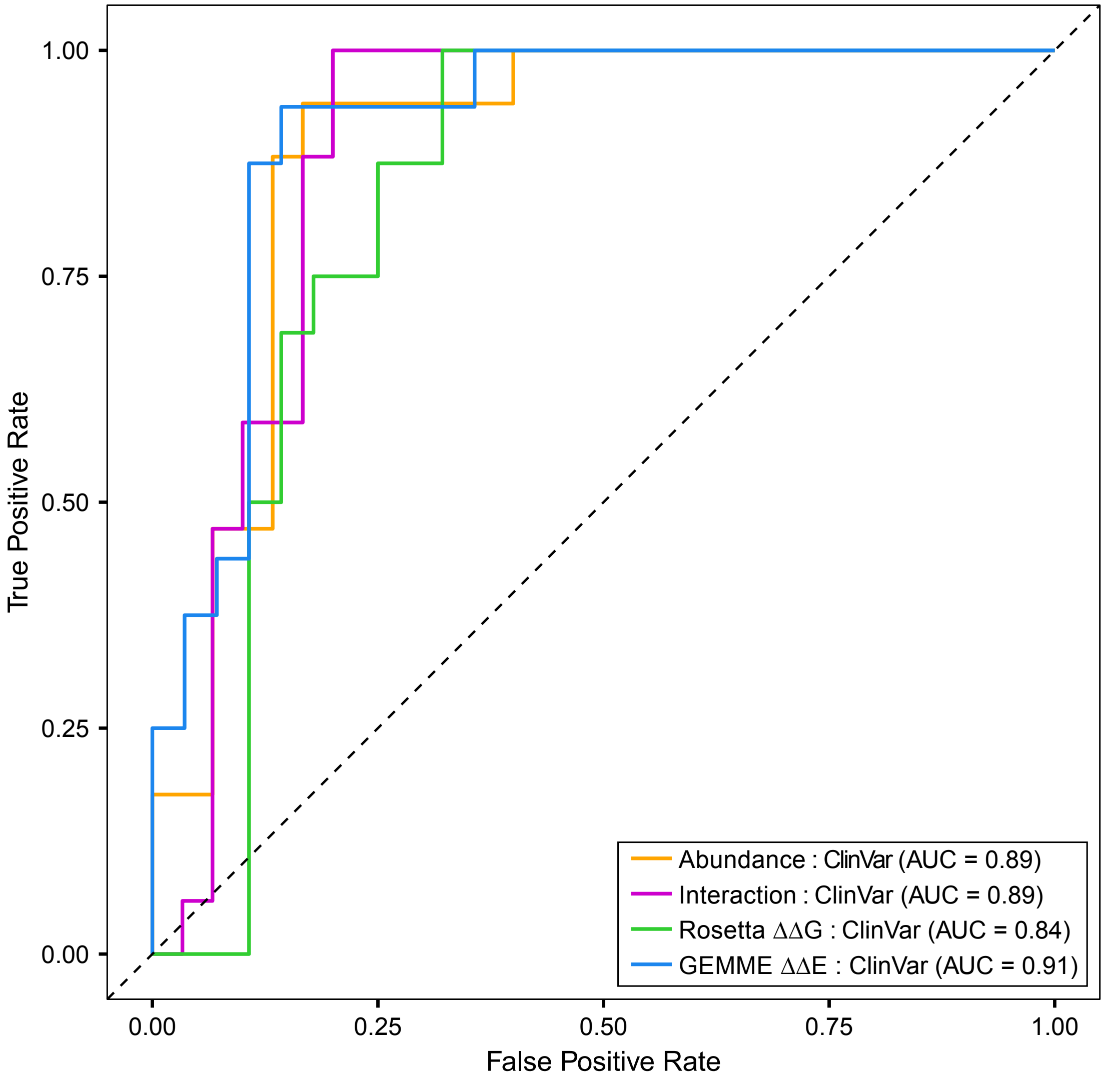


**Fig. S14** – *Classification of MLH1 pathogenicity using experimental and computational scores*. ROC curves to assess how well our experimental abundance and interaction scores, as well as computational Rosetta and GEMME predictions perform in separating pathogenic (and likely pathogenic) from benign (and likely benign) MLH1 variants. The AUC for each predictor is reported in the legend. The dashed diagonal line denotes a random classifier.

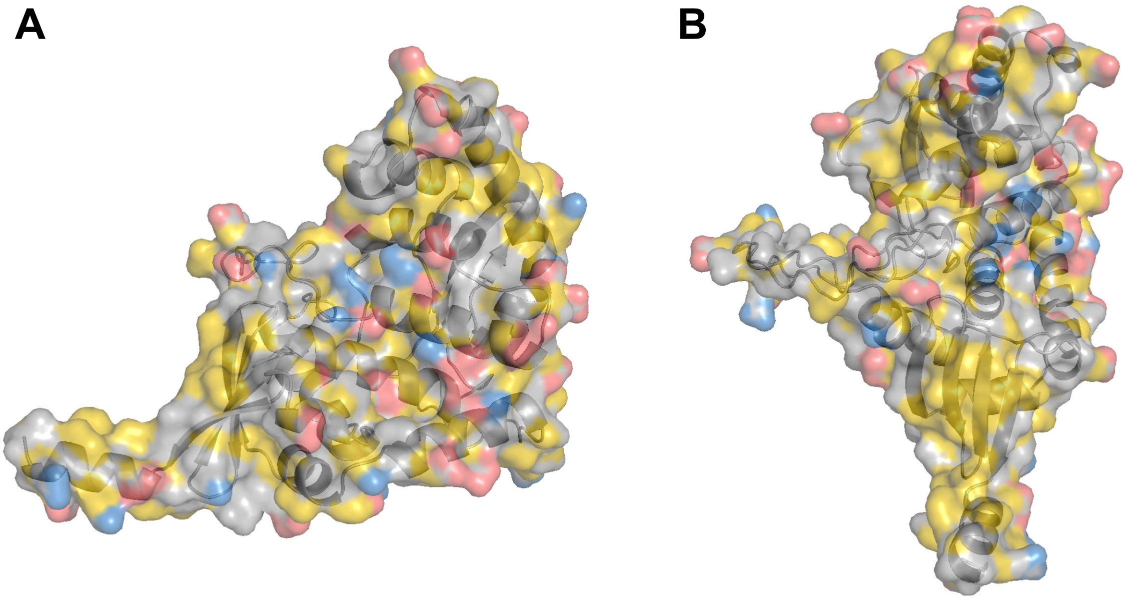


**Fig. S15** – *Hydrophobicity of the canonical PMS2 interface.* (A) Cartoon representation of the C-terminal domain of MLH1 predicted by AlphaFold2. Surface residues are colored according to their chemical properties using the YRB script (see Materials and Methods). Carbon atoms not bound to oxygen or nitrogen atoms are colored yellow (hydrophobic amino acids), negatively charged oxygen atoms are colored red (negatively charged amino acids), positively charged nitrogen atoms are colored blue (positively charged amino acids) and all other atoms are colored gray. The four β-strands in the canonical interface with PMS2 are enriched with hydrophobic amino acids (yellow) and depleted in charged residues (red and blue). (B) Same as in (A), but a different view of the structure highlighting the four β-strands at the bottom front.

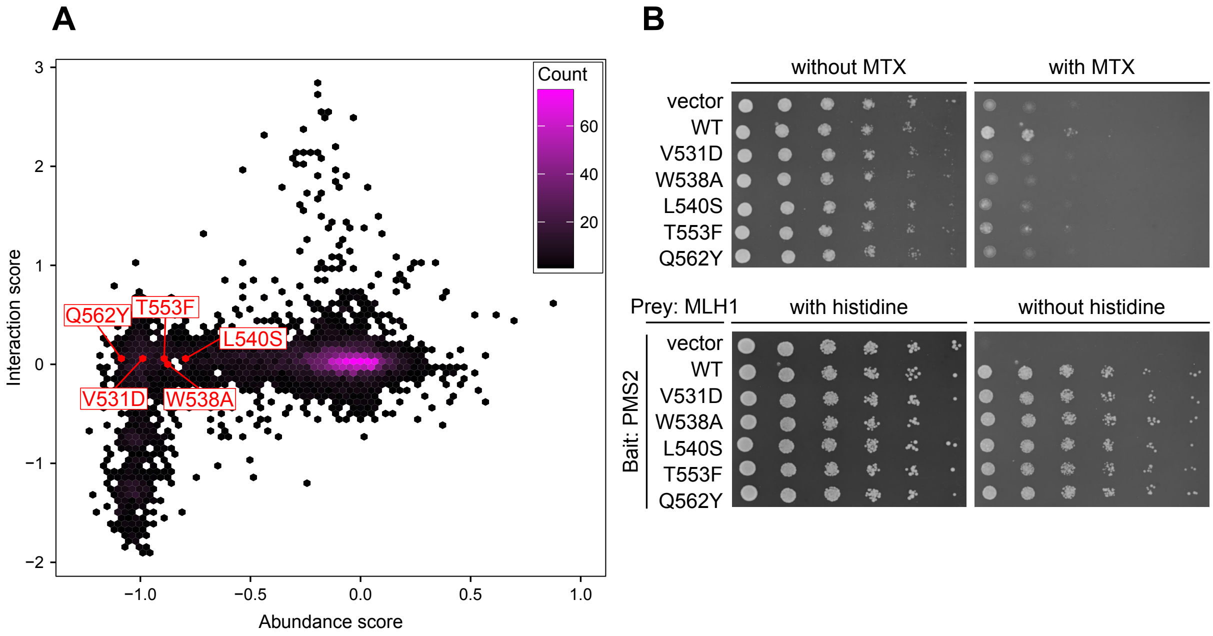


**Fig. S16** – *Dynamic range and sensitivity of the DHFR-PCA and Y2H assay.* (A) Scatter density plot comparing the interaction scores and the abundance scores. The color scale is indicated in the legend. Selected MLH1 variants that displayed reduced abundance but wild-type-like interaction with PMS2 are highlighted in red. (B) Yeast growth assays comparing the growth of a vector, wild-type or the indicated MLH1 variant in the DHFR-PCA (top panel) and the Y2H assay (bottom panel). In the DHFR-PCA, cells were grown on medium with or without MTX. In the Y2H assay, cells were grown on medium with or without histidine.

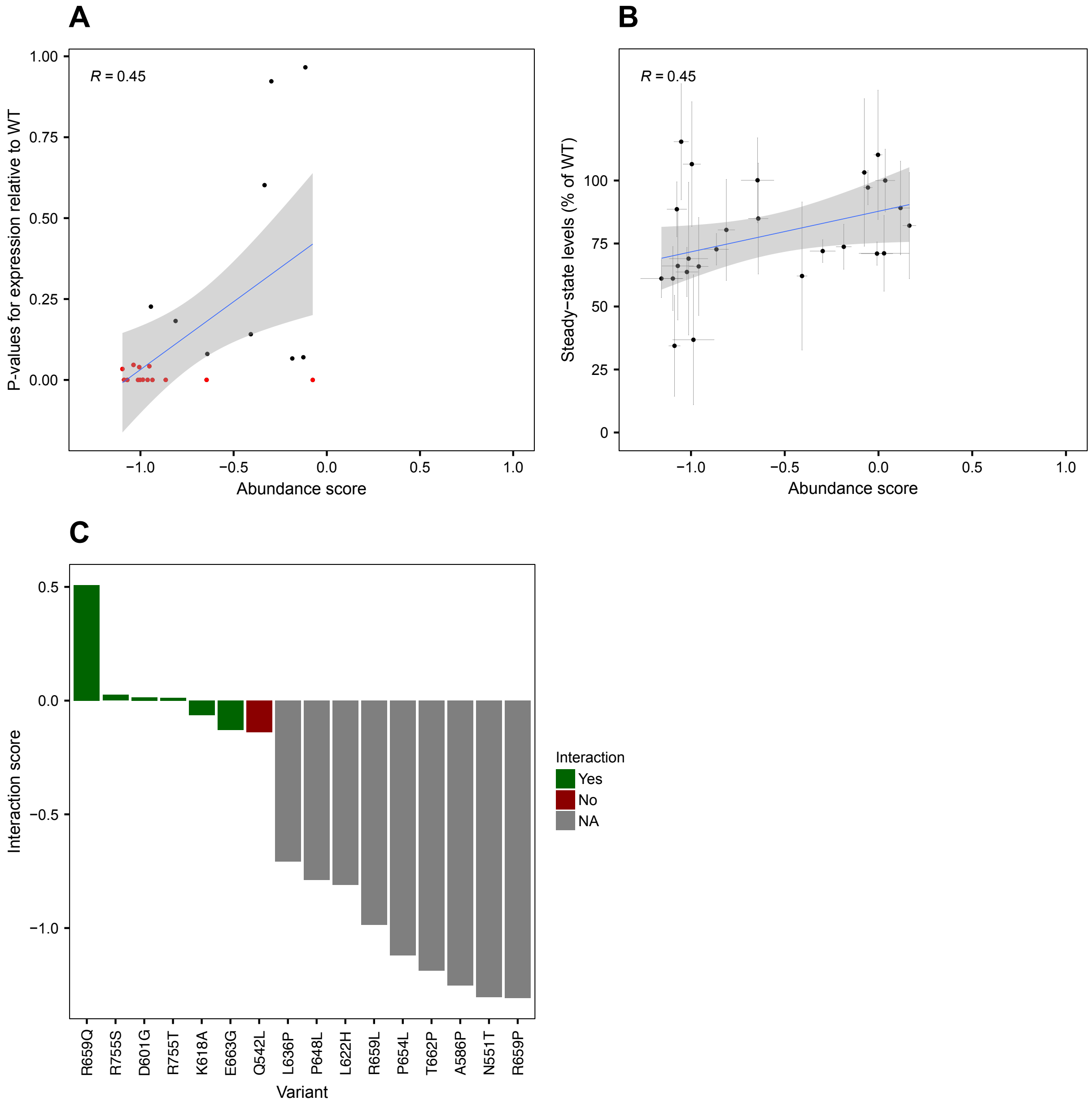


**Fig. S17** – *Benchmarking abundance and interaction scores against existing literature.* (A) Comparison of our experimental abundance scores with protein expression levels reported in [63]. In their study, HEK293T cells were transiently transfected with an MLH1 variant, and cellular extracts were analyzed using SDS-PAGE and immunoblotting with an anti-MLH1 antibody. Statistical analysis of MLH1 expression levels relative to WT was performed using a t-test, with p-values calculated for each variant. Variants shown in red indicate $p < 0.05$. As expected, most of the variants with significantly reduced expression levels were also measured to have reduced abundance scores in our work. (B) Comparison of our experimental abundance scores with protein expression levels reported in [31]. In their study, HCT116 cells were transiently transfected with an MLH1 variant and analyzed using immunofluorescence microscopy with an anti-MLH1 antibody. Expression levels were quantified and reported as a percentage relative to WT. As expected, most of the variants with reduced expression levels were also measured to have reduced abundance scores in our work. (C) Comparison of our experimental interaction scores with PMS2 interaction data reported in [44]. In their study,

HEK293T cells were transiently co-transfected with an MLH1 variant and PMS2, and cellular extracts were analyzed using SDS-PAGE and immunoblotting with anti-MLH1 and anti-PMS2 antibodies. Based on PMS2 expression levels, MLH1 variants were classified as either interacting or not interacting with PMS2. Variants with expression levels too low to measure PMS2 expression were denoted as NA. As expected, most MLH1 variants that interacted with PMS2 in their study also exhibited positive interaction scores or scores close to 0 in our work. Additionally, variants reported in [44] to have dramatically reduced expression levels also displayed reduced interaction scores in our work.
